# Discovery and whole genome sequencing of a human clinical isolate of the novel species *Klebsiella quasivariicola* sp. nov

**DOI:** 10.1101/176743

**Authors:** S. Wesley Long, Sarah E. Linson, Matthew Ojeda Saavedra, Concepcion Cantu, James J. Davis, Thomas Brettin, Randall J. Olsen

**Affiliations:** Center for Molecular and Translational Human Infectious Diseases Research, Department of Pathology and Genomic Medicine, Houston Methodist Research Institute and Houston Methodist Hospital, Houston, Texas, USA; Department of Pathology and Laboratory Medicine, Weill Cornell Medical College, New York, New York USA; Computing, Environment and Life Sciences, Argonne National Laboratory, Argonne IL, 60439, USA; Computation Institute, University of Chicago, Chicago, Illinois, 60637, USA

## Abstract

Originally thought to be a single species, *Klebsiella pneumoniae* has been divided into three distinct species: *K. pneumoniae*, *K. quasipneumoniae* and *K. variicola*. In a recent study of 1,777 extended-spectrum beta-lactamase (ESBL)-producing *Klebsiella* strains recovered from human infections in Houston, we discovered one strain (KPN1705) causing a wound infection that was phylogenetically distinct from all currently recognized *Klebsiella* species. Whole genome sequencing of strain KPN1705 revealed that it was single locus variant of the multilocus sequence type ST-1155. This sequence type was reported only once previously. To further investigate the phylogeny of these two organisms, we sequenced the genome of strain KPN1705 to closure and compared its genetic features to *Klebsiella* reference strains. Results demonstrated strain KPN1705 extensively shares core gene content, antimicrobial resistance genes, and plasmids with *K. pneumoniae*, *K. quasipneumoniae* and *K. variicola*. Since strain KPN1705 and the previously reported novel strain are phylogenetically most closely related to *K. variicola,* we propose the name *K. quasivariicola* sp. nov.

## IMPORTANCE

*K. pneumoniae, K. quasipnuemoniae* and *K. variicola* are serious human pathogens that are increasingly associated with multidrug resistance and high morbidity and mortality. In a recent study of a large, comprehensive, population-based collection of antibiotic resistant *Klebsiella* isolates recovered from human patients, we discovered a novel species that is related to but distinct from *K. variicola*. This clonal group has been reported only once previously. We sequenced the genome of this clinical isolate and compared its genetic features to other *Klebsiella* strains. We propose the name *K. quasivariicola* sp. nov. for this new species.

## OBSERVATION

Members of the genus *Klebsiella* are a common cause of human morbidity and mortality (1, 2). Many community-acquired and healthcare-associated outbreaks of invasive *K. pneumoniae* disease have been reported (3, 4). Over the past two decades, related *Klebsiella* species have been identified as distinct from *K. pneumoniae* and classified (5-8). In a recent large, comprehensive, population based study of 1,777 extended-spectrum beta-lactamase (ESBL) producing *Klebsiella* strains recovered in our clinical microbiology laboratory, we discovered a unique isolate KPN1705 (9, 10). It was genetically related to, but distinct from, *K. variicola*. We sequenced the genome of this strain, which belongs to a new species herein termed *Klebsiella quasivariicola* sp. nov., to closure and compared its genetic features to other *Klebsiella* reference strains.

## RESULTS

### Whole genome sequencing reveals *Klebsiella quasivariicola* sp. nov., a novel *Klebsiella* pathogenic to humans

In a recent study of 1,777 ESBL-producing *K. pneumoniae* isolates recovered from patients in our health care system, we unexpectedly discovered that 28 strains were phylogenetically allied with *K. variicola* (13 strains) and *K. quasipneumoniae* (15 strains)(10). We identified strain KPN1705 as a distinct outlier in the phylogenetic analysis. It shared a common branch with the *K. variicola,* yet was as distant from the *K. pneumoniae, K. quasipneumoniae,* and *K. variicola* reference genomes as they were from each other (Figure 1).

**Fig 1.**
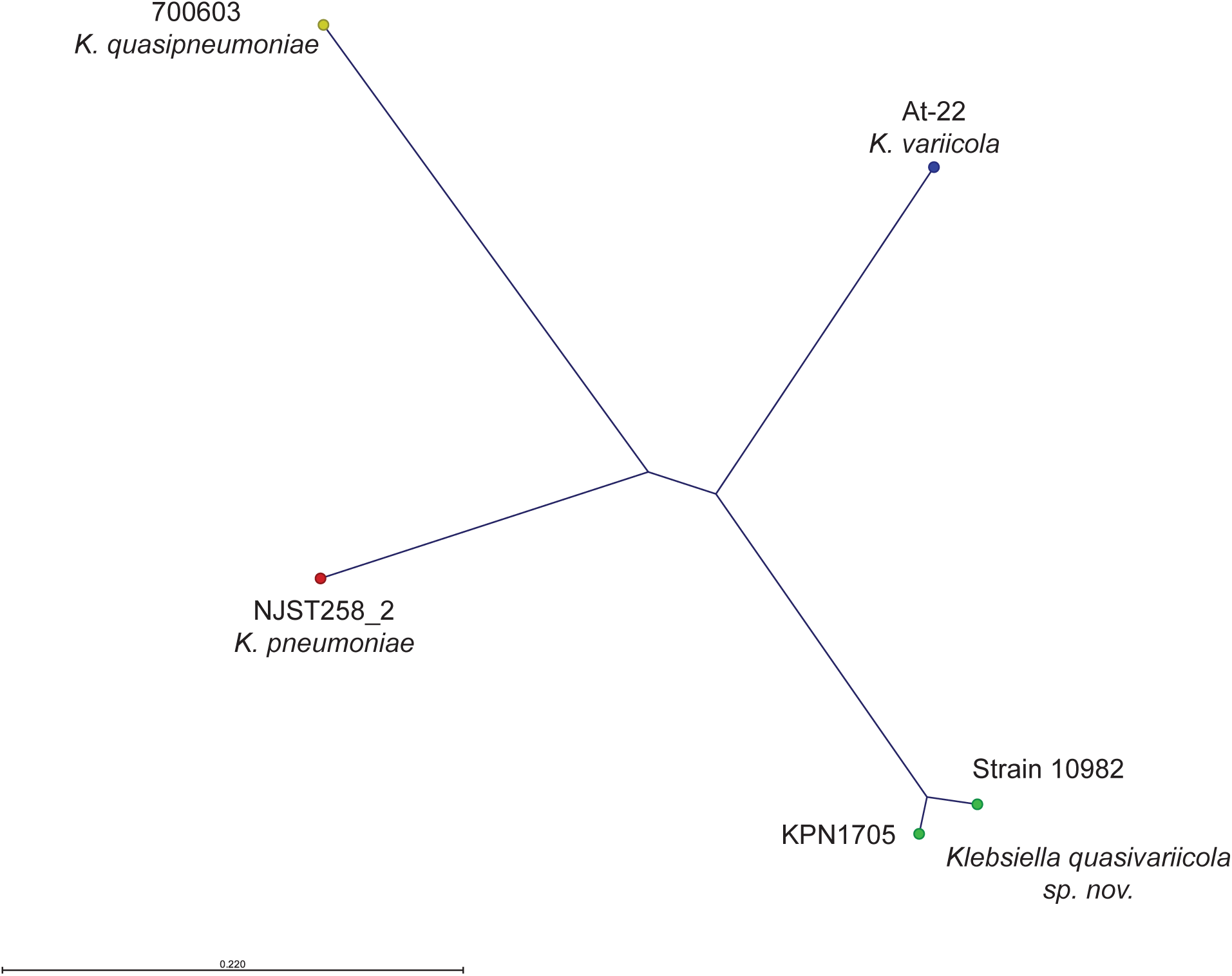
Genetic relationships among *Klebsiella quasivariicola* sp. nov. strains KPN1705 and Strain 10982 and reference strains of *K. pneumoniae* (NJST258_2), *K. variicola* (At-22), and *K. quasipneumoniae* (700603). Phylogenetic relationships were defined by the neighbor-joining method in FastTreeMP with double precision using the closed KPN1705 genome as a reference. The core genome was defined as the chromosomal sequence with the 6 predicted phage sequence regions excluded.

To determine if strains similar to KPN1705 had been previously reported, we determined its multilocus sequence type (MLST). Results revealed that it is a single locus variant of ST-1155, with three SNPs in the *infB*_110 allele. A search of publicly available databases found one previous report of an ST-1155 *Klebsiella*, which was a description of a novel *Klebsiella* dubbed Strain 10982 (11). Strain 10982 was recovered from a perianal swab collected on an ICU patient in Maryland in 2005, as part of a study of AmpC-mediated antimicrobial resistance (11).

To begin assessing the genetic relationship between strain KPN1705 and other *Klebsiella*, we sequenced the genome of KPN1705 to closure. The KPN1705 chromosome is 5,540,188 bp, and three plasmids were identified (described below). Strain 10982 was previously sequenced by Hazen et al. and the assembled 218 contigs are published (11). SNPs were called for reference genomes of *K. pneumoniae* (NJST258_2), *K. quasipneumoniae* (700603), *K. variicola* (At-22), and Strain 10982, using our closed KPN1705 as a reference. The pairwise distance between *K. pneumoniae* and *K. variicola* compared to KPN1705 was 250,000 and 251,939 SNPs, respectively. Similarly, the pairwise distance between *K. pneumoniae* and *K. variicola* and Strain 10982 was 253,227 and 253,864 SNPs. This level of difference between the novel strains and other *Klebsiella* clades is similar to the distance separating the *K. pneumoniae*, *K. variicola* and *K. quasipneumoniae* from one another (mean: 269,799 SNPs, range: 247,050-287,991 SNPs) (Figure 2A). In comparison, KPN1705 and Strain 10982 were closely related, differing from one another by only 34,455 SNPs (Figure 1). This level of difference is similar to the average pairwise distance between any two *K. variicola* strains (average: 38,056 SNPs, range: 31,777 – 45,299) (10). Together, these whole genome sequence data suggest that KPN1705 and Strain 10982 represent a novel *Klebsiella* species, and we propose the name *Klebsiella quasivariicola* sp. nov.

**Fig 2.**
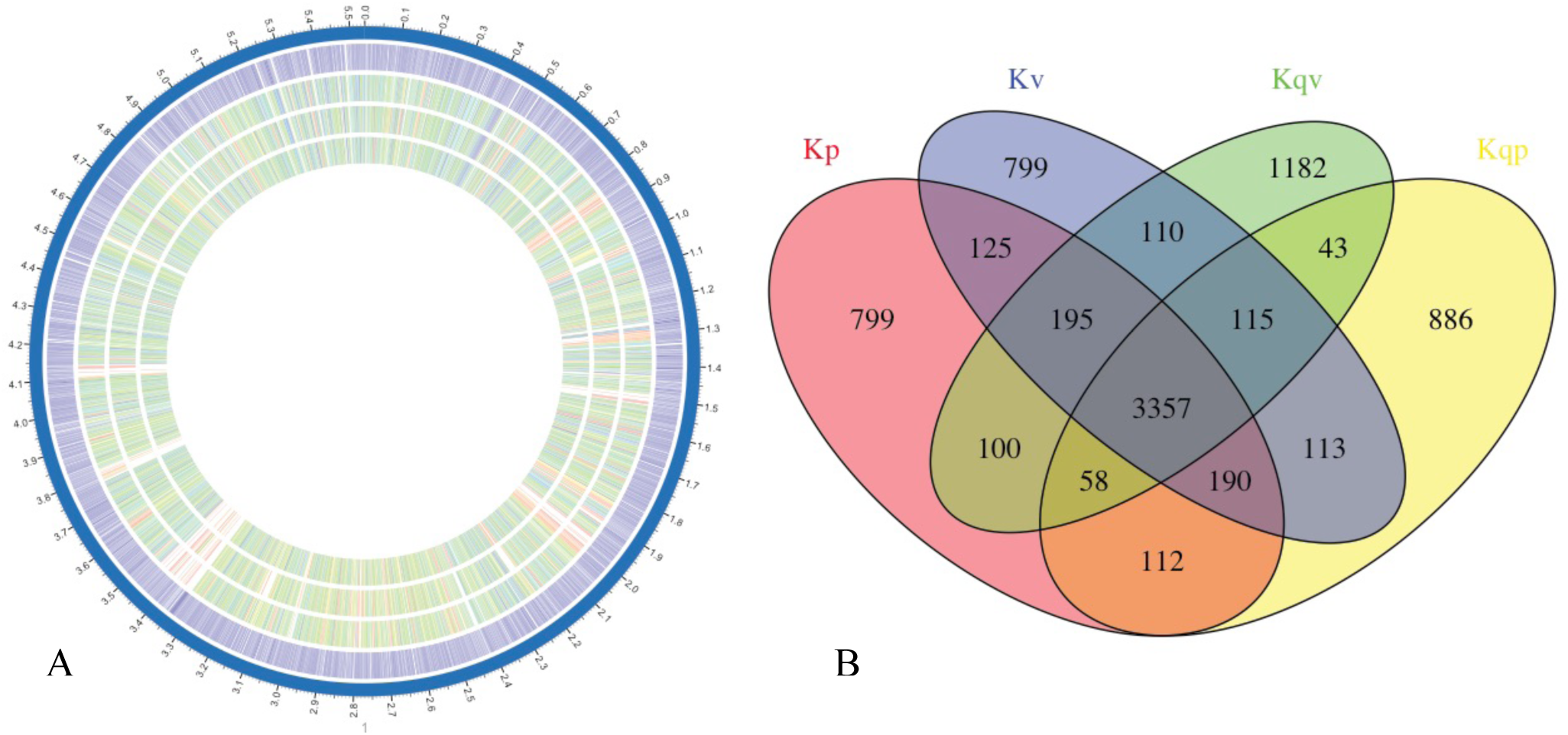
Gene content differences between the reference genomes *Klebsiella quasivariicola* sp. nov. (KPN1705), *K. pneumoniae* (NJST258_2), *K. variicola* (At-22), and *K. quasipneumoniae* (700603). A. Bidirectional BLAST was performed by using the PATRIC resource to illustrate the differences in gene content between these two reference genomes. The color indicates the percent identity of the BLAST hit for each gene, with darker shading indicating a bidirectional hit and lighter shading indicating a unidirectional hit. Outer-most ring is the KPN1705 chromosome reference, followed by *K. pneumoniae* NJST258_2, *K. quasipneumoniae* 700603, and *K. variicola* At-22 on the innermost ring. B. Venn diagram showing shared gene content between the *K. pneumoniae* NJST258_2 (Kp, red), *K. quasipneumoniae* 700603 (Kqp, yellow), *K. variicola* At-22 (Kv, blue), and *K. quasivariicola* sp. nov. (Kqv, green) as determined by Roary.

### Plasmid and phage content in *Klebsiella quasivariicola* sp. nov. strain KPN1705

Next, we characterized the plasmids carried by strain KPN1705. Using our assembled whole genome data, we identified three plasmids, pKPN1705-1 (240,771bp), pKPN1705-2 (97,896bp), and pKPN1705-3 (67,851bp). These plasmids were similar to others found in *Klebsiella* species and carried a diverse array of replicons and antimicrobial resistance genes. Six intact phage regions were predicted in the core chromosome, consisting of 359 coding sequences in 322.7 kb of core chromosomal sequence (Table S1 Phage).

### Antimicrobial gene content in *Klebsiella quasivariicola* sp. nov

The SHV-LEN-OKP beta-lactamases are core chromosomal genes of *Klebsiella* that are usually segregated by *Klebsiella* species: *K. pneumoniae* (SHV restricted), *K. quasipneumoniae* (OKP restricted), and *K. variicola* (LEN restricted) (8, 12, 13). SHV beta-lactamase genes can also be carried on plasmids (14). We assessed the antimicrobial gene content of KPN1705 and determined it carries the LEN-24 beta-lactamase on its chromosome, similar to what is commonly found in *K. variicola*. This further contributed to our suggestion to call this novel species *K. quasivariicola* sp. nov. KPN1705 also carried the gene encoding the SHV-30 ESBL enzyme on plasmid pKPN1705-3. Genes encoding KPC, OXA, CTX-M, TEM and NDM-1 were not detected.

### Gene content comparison between *K. pneumoniae, K. variicola, K. quasipneumoniae* and *K. quasivariicola*

We compared the gene content between our ESBL-producing *K. pneumoniae, K. variicola*, *K. quasipneumoniae* and *K. quasivariicola* sp. nov.. We identified a total of 8,184 unique genes present in the pangenome of all four species (Figure 2B). A *Klebsiella* core genome consisted of 3,357 unique genes that were present in the reference genome of each clade. A bidirectional BLAST comparing the 4 reference genomes to KPN1705 shows the distance between each is similar, with gaps present in the regions corresponding to 6 predicted phage regions (Figure 2A). A table of the gene presence or absence is included in the supplemental (Table S2 Gene Content).

## DISCUSSION

*K. pneumoniae* is a well-known cause of human morbidity and mortality. Although less common, the closely related organisms *K. variicola* and *K. quasipneumoniae* also cause life-threatening infections (5, 7, 10, 15). The difficulty that conventional clinical microbiology laboratories have in distinguishing *K. variicola* and *K. quasipneumoniae* from *K. pneumoniae* may contribute to our underestimation of their potential as human pathogens (10, 16). The discovery of this novel clade of *Klebsiella,* the *K. quasivariicola* sp. nov., represents yet another *Klebsiella* species capable of causing serious human infections. Importantly, when novel strain 10982 was first described, the investigators questioned whether it had simply colonized the gastrointestinal tract or if it was potentially pathogenic. Our novel strain KPN1705 was recovered from a wound culture, strongly suggesting a causative role for the abscess. In addition, the detection of multiple antimicrobial resistance genes, including a SHV ESBL enzyme, increases its virulence potential.

Our whole genome sequence data provides clues to the relationships between the *Klebsiella* clades. The core genome content of *K. quasivariicola* sp. nov., is similar to *K. pneumoniae*, *K. variicola* and *K. quasipneumoniae*, despite the extensive diversity that has been reported to occur within and between clades (6, 9, 10). Also, consistent with previous reports, (6) we observed the plasmids present in KPN1705 to be similar to those found in other *Klebsiella* species. Importantly, these plasmids carry multiple genes encoding virulence factors and antimicrobial resistance genes (17).

These data provide new insight to the natural history and pathogenesis of *Klebsiella* organisms. Additional strains of *Klebsiella quasivariicola* sp. nov. are needed to better characterize this new species. Improved diagnostic methods or widespread use of whole genome sequencing of clinical isolates may be necessary to ensure timely and appropriate identification of these pathogens.

## MATERIALS AND METHODS

### Whole genome sequencing of *Klebsiella*

The genome of strain KPN1705 was previously described using Illumina short read data (9). To obtain long reads to close the genome, we sequenced the genome of strain KPN1705 to closure using the 1D Ligation sequencing kit, R9.4 flow cell, and Oxford Nanopore Technologies MinION Mk-Ib sequencer.

### Bioinformatics analysis of strains

The single nucleotide polymorphism calling pipeline and additional bioinformatics pipelines were described previously (9). BLAST was performed using the NCBI BLAST toolkit and CLC Genomics Workbench v.10.1. Visualization of SNP distribution was performed using CLC Genomics Workbench v.10.1. FASTQ files were assembled into contigs using Spades v3.10.1, and contigs were annotated using Prokka v1.12 (18, 19). Unicycler v0.4.0 was used for hybrid assembly and polishing of short reads and long reads into a closed genome for KPN1705 (20). Gene content analysis was performed using Roary v3.6.1 (21). Bidirectional BLAST and circos visualization were performed using PATRIC (www.patricbrc.org). Assembly of SNPs into phylogenetic trees was accomplished with the scripts prephix v3.3.0, phrecon v4.6.0, and FastTreeMP v2.1 (22). Phage regions were predicted using PHASTER (23). Prephix and phrecon are available from https://github.com/codinghedgehog. The Venn diagram was made in RStudio 1.0.136 using R 3.3.2 and the VennDiagram package v1.6.17 (24).

### MALDI-TOF Identification

KPN1705 was isolated by the Houston Methodist Diagnostic Microbiology Laboratory as described previously (25).

### Accession Numbers

The genomes of the strain sequenced for this study have been deposited in the NCBI database under BioProject PRJNA376414 and BioSample SAMN06438648. The accession numbers for the KPN1705 closed genome and plasmids are CP022823-CP022826. Reference genome Genbank accession numbers are as follows: NJST258_2 (CP006918.1), 700603 (CP014696.2), At-22 (CP001891.1), and Strain 10982 (GCA_000523395.1).

## ACKNOWLEDGEMENTS

This work was supported by funds from the National Institute of Allergy and Infectious Diseases, National Institutes of Health, Department of Health and Human Service (contract HHSN272201400027C). We acknowledge Dr. J. Kristie Johnson for helpful discussions regarding Strain 10982, and Kathryn Stockbauer for help in preparing this manuscript.

